# Co-release of GABA and ACh from medial olivocochlear neurons fine tunes cochlear efferent inhibition

**DOI:** 10.1101/2024.08.12.607644

**Authors:** Tais Castagnola, Valeria C Castagna, Siân R. Kitcher, Lester Torres Cadenas, Mariano N Di Guilmi, Maria Eugenia Gomez Casati, Paula I Buonfiglio, Viviana Dalamón, Eleonora Katz, Ana Belén Elgoyhen, Catherine J.C. Weisz, Juan D Goutman, Carolina Wedemeyer

## Abstract

During development, inner hair cells (IHCs) in the mammalian cochlea are unresponsive to acoustic stimuli but instead exhibit spontaneous activity. During this same period, neurons originating from the medial olivocochlear complex (MOC) transiently innervate IHCs, regulating their firing pattern which is crucial for the correct development of the auditory pathway. Although the MOC-IHC is a cholinergic synapse, previous evidence indicates the widespread presence of gamma-aminobutyric acid (GABA) signaling markers, including presynaptic GABA_B_ receptors (GABA_B_R). In this study, we explore the source of GABA by optogenetically activating either cholinergic or GABAergic fibers. The optogenetic stimulation of MOC terminals from GAD;ChR2-eYFP and ChAT;ChR2-eYFP mice evoked synaptic currents in IHCs that were blocked by α-bungarotoxin. This suggests that GABAergic fibers release ACh and activate α9α10 nicotinic acetylcholine receptors (nAChRs). Additionally, MOC cholinergic fibers release not only ACh but also GABA, as the effect of GABA on ACh response amplitude was prevented by applying the GABA_B_-R blocker (CGP 36216). Using optical neurotransmitter detection and calcium imaging techniques, we examined the extent of GABAergic modulation at the single synapse level. Our findings suggest heterogeneity in GABA modulation, as only 15 out of 31 recorded synaptic sites were modulated by applying the GABA_B_R specific antagonist, CGP (100-200 µM). In conclusion, we provide compelling evidence that GABA and ACh are co-released from at least a subset of MOC terminals. In this circuit, GABA functions as a negative feedback mechanism, locally regulating the extent of cholinergic inhibition at certain efferent-IHC synapses during an immature stage.

**Significance statement:** Before hearing onset, the medial olivocochlear (MOC) efferent system of the mammalian cochlea regulates the pattern of IHC spontaneous firing rate through the activation of α9α10 nAChRs. However, GABA is also known to have a modulatory role at the MOC-IHC synapse. Our results show that GABA is co-released from at least a subset of MOC terminals, working as a precise regulatory mechanism for ACh release. Furthermore, we demonstrate that not all synaptic contacts within a single IHC are equally modulated by GABA.

## Introduction

Co-release of different neurotransmitters from the same neuron was described many years ago (Hnasko & Edwards, 2012). Recent findings suggest that several populations of neurons in the central nervous system (CNS) previously thought to release only glutamate, acetylcholine, dopamine, or histamine, also release GABA (reviewed in (Tritsch et al., 2016). Consistent with this, extensive co-expression of GABAergic and cholinergic markers is both widespread and evolutionarily conserved in vertebrate species (O’Malley and Masland, 1989; Lee et al., 2010; Granger et al., 2016, 2020), suggesting that GABA and ACh are co-released from some cholinergic neurons in the CNS. Given that GABA acts through both ionotropic and metabotropic receptors localized either pre and/or postsynaptically (Eccles et al., 1954; Turecek and Trussell, 2001; Magnusson et al., 2008; Pugh and Jahr, 2011; Wedemeyer et al., 2013; Clause et al., 2014; Zorrilla de San Martin et al., 2017) co-release could be an important means of modulating and fine-tuning synaptic transmission.

In the mammalian inner ear, sound is converted into electrical signals by inner and outer hair cells (IHCs and OHCs, respectively). These signals are sent to the CNS mainly via Type I spiral ganglion afferent neurons that contact the IHCs (Spoendlin and Schrott, 1988). However, during development, IHCs do not respond to acoustic stimuli but instead exhibit intrinsically generated electrical activity (Tritsch et al., 2007, 2011; Sendin et al., 2014).Within this same period, medial olivocochlear (MOC) neurons located in the ventral brainstem (VNTB) send efferent projections to the IHCs (Guinan et al., 1996; Guinan, 2011). This cholinergic and inhibitory innervation plays a significant role in modulating developing IHCs excitability (Glowatzki and Fuchs, 2000; Simmons, 2002; Goutman et al., 2005) and then disappears by the onset of hearing (postnatal day (P) 12-14 in altricial rodents) (Simmons, 2002; Katz, 2004; Roux et al., 2011). Tight regulation of this prehearing activity by MOC innervation is essential for the precise development and refinement of the auditory pathway (Galambos, 1956; Johnson et al., 2013; Clause et al., 2014, 2017; Di Guilmi et al., 2019; Wang et al., 2021).

The main transmitter at MOC-hair cell synapses is acetylcholine (ACh). This neurotransmitter activates calcium-permeable α9α10 nicotinic ACh receptors (nAChRs; Elgoyhen et al., 1994, 2001) that are functionally coupled to calcium-dependent SK and/or BK potassium channels that ultimately hyperpolarize the cells (Dulon and Lenoir, 1996; Glowatzki and Fuchs, 2000; Oliver et al., 2000; Katz et al., 2004; Wersinger et al., 2010; Wersinger and Fuchs, 2011). It has been shown that this synapse is subject to modulation from different sources. Thus, glutamate released from IHCs enhances the release of ACh and subsequently potentiates MOC inhibition through a negative feedback mechanism mediated by metabotropic glutamate receptors (mGlu1) (Ye et al., 2017) . In addition, nitric oxide produced by IHCs (Kong et al., 2013) and ACh through presynaptic nicotinic acetylcholine receptors (Zhang et al., 2020) increases the ACh release probability from MOC neurons.

Abundant GABAergic markers have been found below the IHC and OHC areas (Fex and Altschuler, 1986; Vetter et al., 1991; Eybalin, 1993; Maison et al., 2003; Bachman et al., 2024). In adult mice, GABA co-localizes with ACh at almost all synapses of the OC system (Maison et al., 2003). Furthermore, during postnatal development before hearing onset when MOC fibers transiently innervate the IHCs, ACh release at this synapse is downregulated by GABA acting on presynaptic GABA_B_R (Wedemeyer et al., 2013). Even though this previous work indicated a clear role for GABA at the MOC-IHC synapse, the question remained as to whether GABAergic efferent fibers are a subpopulation of the MOC fibers or if the same cholinergic fibers also express GABA and co-release this neurotransmitter together with ACh.

In the present work we aimed to answer this question by using a combination of optogenetic, immuno-localization techniques and optical GABA detection experiments. Our results suggest that during development, ACh and GABA are co-released from a subset of MOC terminals innervating the IHCs. In addition, using calcium imaging techniques at single synapses, we show that not all of the multiple synapses that reach a single IHC are sensitive to the GABA_B_R modulation, indicating a high degree of heterogeneity of neurotransmission during this transient developmental innervation.

## Materials and Methods

### Animal procedures

Euthanasia and tissue extraction were carried out according to approved animal protocols (INGEBI and The International Guiding Principles for Biomedical Research Involving Animals and The National Institutes of Health guidelines, NIH-OLAW OMB Number 0925-0765). Male and female mice were used in experiments. Mouse lines included wildtype (WT) Balb/C, ChAT-Cre (Jax Cat No: 006410), GAD-CreERT2 (Jax Cat No: 010702), Ai14 tdTomato reporter mice (Jax Cat No: 007914), and ChR2-eYFP (Jax Cat No: 012569). Either double homozygous or hemizygous mice for ChAT-Cre and homozygous for ChR2-eYFP animals were used, hereafter referred to as ChAT;ChR2-eYFP. Special care was taken not to include any animal with a possible ectopic expression (Chen et al., 2018). For optogenetic activation of GABAergic fibers, GAD-CreERT2 animals were crossed with ChR2-eYFP mice (referred to as GAD;ChR2-eYFP). To induce the expression of ChR2, mice were intraperitoneally injected with 100 mg per gram of body weight of tamoxifen (Sigma-Aldrich T5648) diluted in sesame oil (Sigma-Aldrich Cat No: S3547) during 4 consecutive days, starting at P3. Littermates hemizygous for GAD-CreERT2 and homozygous for ChR2 were detected with PCR and used in the experiments. For the immunohistochemistry assays, a tdTomato reporter mouse line was crossed either to GAD-CreERT2 (GAD;tdTomato) or ChAT-Cre mice (ChAT;tdTomato), and PCR was performed to detect hemizygous offsprings for both transgenes. For fluorescence imaging of iGABABSnFr, Ngn-CreERT2 (Jax Cat No: 008529) and Bhlhb5-Cre mice (on a Sv/129, C57BL/6J mixed background, MGI No:4440795) were used. Mice were housed on a 12/12 hr light/dark cycle, with continuous availability of food and water.

### Electrophysiological recordings from IHCs

For patch-clamp recordings from IHCs, apical turns of P9-11 mice cochleae were dissected and placed under an insect pin attached to a round glass coverslip with Sylgard (Dow Chemicals, Midland, MI, USA). Only one cell was recorded per cochlea. Tissue was initially visualized using an upright Zeiss Axioscope microscope with a sCMOS Zyla camera (Andor Technology Ltd) or Olympus BX51WI microscope (Olympus Corporation) with an EM-CCD camera (Andor iXon 885, Andor Technology Ltd). Recordings were then performed using 40X or 60X water immersion objectives with DIC optics.

For IHC recordings, evoked IPSCs were obtained in the whole-cell voltage clamp configuration with optogenetic or extracellular electrical stimulation of the MOC fibers. The holding potential (V_h_) was maintained at -80 mV for optogenetics experiments. In calcium imaging experiments, to maximize Ca^2+^ driving force, IHCs were voltage clamped at -120 mV during a brief period (500 ms) around MOC stimulation which coincided with the fast imaging interval. Otherwise, IHCs were clamped at -70 mV. Membrane voltages were not adjusted for the liquid junction potential (-4 mV). Patch-clamp recordings were performed using 1 mm diameter borosilicate glass micropipettes (World Precision Instrument, Cat No: 1B100F-4) with tip resistances of 6-7.5 MΩ. Recordings were performed in voltage-clamp using an Axopatch 200B (Molecular Devices) or Multiclamp 700B (Molecular Devices) amplifier with 1440A Digidata or BCN 2120 board (National Instruments). Recordings were sampled at 10-50 kHz and lowpass filtered at 2-10 kHz. Data was acquired with WinWCP software (J. Dempster, University of Strathclyde) or pClamp 9.2 (Molecular Devices). All recordings were analyzed with custom-written routines in IgorPro 6.37 (Wavemetrics). IPSCs were identified automatically with a search routine based on an amplitude threshold (>3 SD of the baseline noise) and an integral of the event trace threshold (Moglie et al., 2018).

The cochlear preparation was superfused continuously at room temperature and at a rate of ∼ 2–3 ml/min with extracellular saline solution of an ionic composition similar to that of the cochlear perilymph (in mM): 155 NaCl, 5.8 KCl, 1.3 CaCl_2_, 0.9 MgCl_2_, 0.7 NaH_2_PO_4_, 5.6 D-glucose, 10 HEPES buffer, pH 7.4, osmolarity ∼315 mOsm. For iGABASnFR experiments, extracellular solution was the same, but for zero calcium experiments, CaCl_2_ was substituted with MgCl_2_ to maintain equal total divalent concentration, and 1 mM EGTA was added. To increase the depolarization of efferent terminals in GAD;ChR2-eYFP experiments, the calcium concentration was increased to 1.8 mM and 4-aminopyridine (2 mM) was added in the solution. Internal solution contained (in mM): 140 KCl, 3.5 MgCl_2_, 0.1 CaCl_2_, 5 EGTA, 5 HEPES,2.5 Na2ATP. For calcium imaging experiments recording pipettes were filled with an internal solution of the following composition (in mM): 95 KCl, 40 K-ascorbate, 5 HEPES, 2 pyruvate, 6 MgCl_2_, 5 Na2ATP, 10 phosphocreatine-Na_2_, 0.5 EGTA, 0.4 Fluo-4 (calcium indicator). In all cases pH was adjusted to 7.2 with KOH and osmolarity was ∼ 290 mOsm. Series resistance was not compensated for.

Drug application was performed via addition to the re-circulating bath solution (5-10 min). Drugs were obtained from Sigma-Aldrich, Inc. (St. Louis, MO, USA), Alomone Labs (Jerusalem, Israel), Tocris Bioscience and Thermo Fisher Scientific (Waltham, MA, USA).

### Electrical and optogenetic stimulation of MOC axons

Electrically evoked inhibitory postsynaptic currents (*e*IPSCs) were generated through unipolar electrical stimulation of the MOC efferent terminals as described previously (Goutman et al., 2005; Zorrilla de San Martin et al., 2010). Briefly, the electrical stimulus was delivered via a 20-to 80- μm-diameter glass pipette placed at 20- 50 μm modiolar to the inner spiral bundle (ISB). To optimize stimulation, cochlear supporting cells were gently removed using a glass pipette with a broken tip. MOC stimulation was performed using an electrically isolated constant current source (model DS3, Digitimer). For calcium imaging experiments stimulation pulses were 100–500 mA, 0.2-2 ms width, 0.2 Hz and for iGABASnFR 50 Hz, 1 second train duration. Stimulation timing and rate was controlled by the PClamp software.

For optogenetically evoked inhibitory postsynaptic currents (*o*IPSCs), 30 blue light pulses of 2 ms duration were elicited at 0.03 Hz (ThorLabs, 480 nm LED light, ∼10.2 mW). The rundown of *o*IPSCs was assessed by repeatedly stimulating cholinergic neurons in ChAT;ChR2-eYFP (2 ms, 0.03 Hz, 480 nm LED, 15 cells, 9 mice), without any drug application, for a maximum of 30 minutes. *o*IPSC amplitudes were measured and normalized to the average of the first 10 light pulses. Data was fitted to a linear regression to estimate the decay of the amplitude of the response over time (slope value = 0.01406 pA/s). For those experiments in which GABA_B_-R were blocked, the protocol involved stimulating with 10-20 light pulses while perfusing the preparation with extracellular solution. Then, 200 µM CGP 36216 was applied, and after a 2-minute interval, stimulation was resumed (10-50 pulses). Finally, data was corrected for rundown based on the slope obtained from the linear regression previously described.

### iGABASnFR experiments

#### Posterior semi-circular canal (PSC) AAV injections

Posterior semicircular canal (PSC) injections to introduce AAV particles into the cochlea were performed as described in (Isgrig and Chien, 2018), using aseptic procedures. In brief, neonatal pups (P1-2) were hypothermia anesthetized for ∼5 mins until they did not respond to stimulation, and then remained on an ice pack for the duration of the procedure. A postauricular incision was made using micro-scissors and the skin retracted. The PSC was identified under a surgical microscope and a glass micropipette pulled to a fine point was positioned using a micro-injector (World Precision Instruments). For each mouse only one ear was injected with ∼1.2 µL AAV solution containing gene sequences encoding iGABASnFR variants (iGABASnFR2.0, or FLEX.iGABASnFR2.0). The incision was closed using a drop of surgical glue. About 4-5 pups per litter were injected. Pups were recovered to normal body temperature on a warming pad, while receiving manual stimulation to aid recovery.

#### iGABASnFR imaging

For fluorescence imaging of iGABABSnFR-transduced cells in acutely dissected cochlear preparations, the euthanasia, dissections, extracellular solutions, drug application, and electrical stimulation of MOC axons were as above described for patch-clamp recordings. In zero calcium experiments, calcium chloride was excluded from the extracellular solution and replaced by equimolar magnesium chloride, and 1 mM EGTA was added to further buffer any residual extracellular calcium. iGABASnFR2 and FLEX.iGABASnFR2 were kindly gifted from the laboratory of Dr. Loren Looger and the GENIE Project at Howard Hughes Medical Institute Janelia Research Campus, and then packaged into AAV particles by Signagen.

iGABASnFR expression was targeted to SGN by the PHP.eB AAV serotype and human synapsin (hSynap) promoter. In some iGABASnFR imaging experiments, non-Cre-dependent virus ((PHP.eB)-syn.iGABASnFR2-WPRE) was injected into C57BL/6J mice (7 out of 14 mice). In the remaining 7 mice, Cre-dependent ((PHP.eB)-syn.FLEX.iGABASnFR2-WPRE) virus was injected into the cochlea of Bhlhb5-Cre; tdTomato mice, which results in iGABASnFR expression specifically in Cre-expressing cochlear neurons. In all experiments, the iGABASnFR or tdTomato fluorescence was localized to cells with the clear morphology of type I SGN dendrites, and results were pooled (for analysis see *Image Processing* below). To test whether iGABASnFR fluorescence transients were due to activity-dependent calcium influx that triggers subsequent neurotransmitter-containing vesicle release, in some experiments MOC stimulation-evoked iGABASnFR fluorescence was tested in control conditions, in the zero calcium solution (above) , and again in a return to normal 1.3 mM CaCl2, in the same imaging location.

iGABASnFR imaging was performed on a Nikon A1R upright confocal microscope using resonant scanning in both red (568 nm, for tdTomato imaging in cochlear neurons from transgenic mice) and green (488 nm, for iGABASnFR variants) channels. In some experiments, a DIC-like image was simultaneously collected using the transmitted light detector, which converts the laser signal into a greyscale 3D image. Imaging settings included line averaging of 4-16 lines, bi-directional scanning, 512-1024 resolution, and frame rates of ∼7-15 frames per second.

#### Imaging processing

Fluorescence intensity changes were measured in iGABASnFR-transduced neurons using ImageJ (NIH). A maximum intensity projection of the green (iGABASnFR) image stack was generated and then thresholded to set the regions to be used for region-of-interest (ROI) selection. Thresholds were set to 121, but manually adjusted in the case of tissue with brighter or dimmer fluorescence background (mean = 119 ± 20). The ‘analyze particles’ function was used to automatically draw ROIs around the iGABASnFR-expressing structures of interest, which had the clear morphology of type I SGN. It was not possible to identify individual neurons from either tdTomato or iGABASnFR images, so ROIs likely contain multiple neuron segments. These ROIs were then used to measure fluorescence intensities in the original image stack for each frame. Fluorescence intensity values per ROI and per frame were imported into Origin v2021 (OriginLabs, MA, USA). The baseline mean and standard deviation (SD) of fluorescence was measured for one second prior to electrical MOC axon stimulation. To determine the ΔF/F, first we determined whether each ROI had a positive ‘response’ to the stimulation, defined here as a maximum fluorescence greater than twice the mean of the baseline plus two standard deviations of the baseline (mean + 2SDs). To prevent a noisy fluorescence signal from giving an artificially high ‘maximum’ intensity, we used a rolling average (5 frames before, 5 frames after) to smooth the trace, and determined the fluorescence maximum from this rolling average trace. We then calculated the mean and standard deviation from 1 sec prior to stimulation in the non-averaged trace. The mean + 2SDs was subtracted from the ‘maximum’ to detect positive values that were categorized as a ‘response’. To determine the ΔF/F of the fluorescence following MOC axon stimulation, the mean of the baseline fluorescence was subtracted from the maximum fluorescence, then divided by the mean baseline fluorescence.

#### Calcium imaging experiments

A basal image of the IHC was used to create a donut-shape mask, leaving the center of the cell body out of the analysis. The mask was then divided into 24 radial ROIs and ΔF was measured in each ROI for each time frame. Photobleaching was corrected by fitting a line between the pre-stimulus baseline and final fluorescence. We considered that there was a significant increase in fluorescence in those cases where the peak fluorescence signal detected after electrical stimulation was threefold higher than the SD of the baseline and the integral of the fluorescence signal was above 0.3 (arbitrary units × seconds). Finally, those ROIs that exhibited a consistent pattern of activation were selected as hotspots. When the increase in fluorescence did not fulfill any of those criteria, the event was considered a synaptic failure. Synaptic failures were counted in each hotspot along the duration of the experiment, making it possible to estimate the probability of detecting a calcium event as:

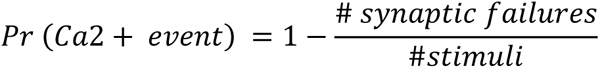

The analysis of the effect of CGP on the activity of individual calcium hotspots required the determination of an amplitude threshold in the calcium signal that took into consideration both the variability and the fluorescence bleaching. As the effect of CGP took approximately 5 minutes to develop completely, control experiments were undertaken in which cells were imaged repetitively while MOC fibers were stimulated, and synaptic parameters were calculated at t = 5 minutes and compared to t = 0. These control experiments produced a mean of 0.7 arbitrary units (A.U) and a standard deviation of 0.6 A.U relative to measurements at t = 0. From this estimate, a z-score threshold = 3 was determined such that experiments that showed a positive deviation compared to control values and exceeded this threshold were considered sensitive to CGP modulation.

#### Immunohistochemistry on brain slices

P9-11 GAD;tdTomato mice were anesthetized via intraperitoneal injection of ketamine (120 mg/kg) and xylazine (25 mg/kg). The animals were then transcardially perfused with 1X phosphate buffered saline (PBS), followed by 4% paraformaldehyde (PFA) in PBS. The brains were postfixed overnight at 4°C in 4% PFA, washed with increasing concentrations of sucrose diluted in PBS (25%, 50% and 75%) and frozen with isopropanol at -70 °C. Thirty micrometer coronal sections were cut using a cryostat (CM1850, Leica). Floating sections were blocked in 10% normal donkey serum (S30-100ML, Millipore) in the case of ChAT;tdTomato mice or 5% normal donkey serum + 5% normal goat serum (S26-100ML, Millipore) for GAD;tdTomato mice, with 1% Triton X-100 in PBS for 2 h at room temperature. The primary antibodies used in this study were as follows: (1) goat anti-choline acetyltransferase (ChAT) (1:300; AB144P, Millipore) to label cholinergic cells; (2) rat anti-Red Fluorescent Protein (RFP) (1:500; 5F8, Chromotek) to label tdTomato positive cells; and (3) mouse anti-glutamic acid decarboxylase (GAD-6) (1:500;MAB351R, Millipore) to label GABAergic cells. The sections were incubated with primary antibodies (diluted in blocking solution) for ∼48 h at 4°C. On those sections where the anti-GAD was used, a second blocking for ∼2 h at room temperature with a Mouse blocker Reagent (2-3 drops in 2.5 ml PBS, MKB-2213-1, Vector Laboratories) was used. The sections were then rinsed in PBS before a 2 h incubation with secondary antibodies at room temperature (1:800; Alexa Fluor 488 donkey anti-goat IgG, 1:800; Alexa Fluor 555 donkey anti-rat IgG,1:800; Alexa Fluor 488 goat anti-mouse IgG2a). After secondary incubation, sections were rinsed and mounted on microscope slides in Vectashield mounting media (Vector Laboratories).

#### Cochlear processing and immunostaining

Cochleae were harvested from P9-11 ChAT;tdTomato mice, as well as from ChAT;ChR2-eYFP and GAD;ChR2-eYFP mice. The tissue was fixed by intra labyrinth perfusion of 4% PFA in PBS and left overnight. After decalcification with 0.12M EDTA, the organ of Corti was microdissected and permeabilized by freeze/thawing in 30% sucrose. The immunohistochemistry procedure followed for mice tissue was the same as indicated earlier for coronal sections, with the exception that in this case the incubation with primary antibody was overnight. Cochleae were blocked in 5% normal goat serum with 1% Triton X-100 in PBS for 2 h, followed by incubation with the primary antibody (diluted in blocking buffer) at 4°C overnight. The primary antibody used was rabbit anti-green fluorescent protein IgG fraction (anti-GFP; 1:2000; A6455, Life Tech). Tissues were rinsed in PBS and incubated with the appropriate secondary antibody (1:1000; Alexa Fluor 488 goat anti rabbit, Invitrogen) for 2 h at room temperature. Finally, tissues were mounted on microscope slides in Vectashield mounting media (Vector Laboratories).

#### Quantification of the Immunostaining

Confocal images were acquired on a Leica TCS SPE Microscope equipped with a 40X and 63X oil-immersion lens. Maximum intensity projections were generated from *z*-stacks and imported to ImageJ software for analysis. JaCoP software was used to calculate the colocalization coefficients (Manders coefficient) of the genetically labeled tdTomato-positive cells (amplified with anti-RFP) with anti-ChAT and anti-GAD antibodies. For Supplementary Figure 3, instead of a z-stack, just one image was chosen in which examples of co-localizing and non co-localizing terminals are seen. Intensity profile lines for both channels were traced along some regions of the inner spiral bundle. In the case of brain slices the bilateral images containing the VNTB were cropped to exclude the lateral superior olive (LSO), where LOC cells reside. Cell counts were performed on monochrome grayscale images of both channels.

### Statistics

All statistical analyses were performed with GraphPad Prism 8.0.2 (GraphPad Software, Inc.). Before performing any analysis, data were tested for normal distribution using the Shapiro–Wilk normality test and parametric or nonparametric tests were applied accordingly. For statistical analyses with two datasets, a two-tailed paired *t*-test or Wilcoxon signed-rank test were used. For comparison between 3 or more conditions Friedman test was used. Finally, a Kolmogorov-Smirnov test was used to analyze different population frequencies. Values of p < 0.05 were considered significant. All data were expressed as mean ± SEM, unless otherwise stated.

## Results

### Optogenetic stimulation of cochlear cholinergic fibers

The conventional approach to evoke neurotransmitter release at MOC-IHC synapses of cochlear explants has primarily involved electrical stimulation or high potassium depolarization of MOC axon terminals (Katz et al., 2004; Gomez-Casati et al., 2005; Goutman et al., 2005; Zorrilla de San Martin et al., 2010; Wedemeyer et al., 2013; Kearney et al., 2019). These studies have provided valuable descriptions of postsynaptic current kinetics, transmitter release properties and both pre-and postsynaptic receptors and ion channels involved in synaptic transmission at this synapse. However, they have the drawback of non-selective activation of adjacent fibers within the stimulated area. Therefore, to take advantage of the ability to stimulate genetically defined cell types, an optogenetic approach was used to activate specific efferent pathways. To validate the use of a transgenic mouse model that expresses Cre recombinase in cholinergic fibers, the expression pattern of the red fluorescent protein tdTomato and ChAT were studied in developing cochleae of the ChAT-Cre mouse line crossed with the Ai14 tdTomato reporter mouse line (Fuchs and Lauer, 2019). Sections from the apical turns of ChAT;tdTomato mouse cochleae (P9-11) were processed and labeled with an anti-ChAT antibody and the tdTomato signal was amplified with anti-RFP antibody. Co-localization of both antibodies was observed in the VNTB where MOC somas are found (**Supplementary Figure 1A-D**). Importantly, there were no cells that were tdTomato+ and anti-ChAT-, indicating that although recombination is not 100%, the ChAT-Cre mouse line is specific for cholinergic neurons. Co-localization of both markers was also found in the inner spiral bundle (ISB) and at the base of OHCs in the outer spiral bundle (OSB) (Manders’ coefficient = 0.81, **Supplementary Figure 1E**), showing that Cre expression follows the previously described innervation pattern of cholinergic efferent neurons in the cochlea (Whitlon and Sobkowicz, 1989; Simmons et al., 1998; reviewed in Simmons, 2002).

Upon establishing that the ChAT promoter drives gene expression in cholinergic fibers within the cochlea, Channelrhodopsin2 (ChR2) was expressed under the control of the ChAT promoter in ChAT-Cre (ChAT;ChR2-eYFP, **Figure 1A**). In these mice, ChR2-eYFP-positive fibers branched extensively in the ISB region and terminated at the base of the IHCs (**Figure 1B**). Fluorescence was absent in hair cells, indicating that the presence of ChR2 was restricted to efferent neurons.

**Figure 1.**
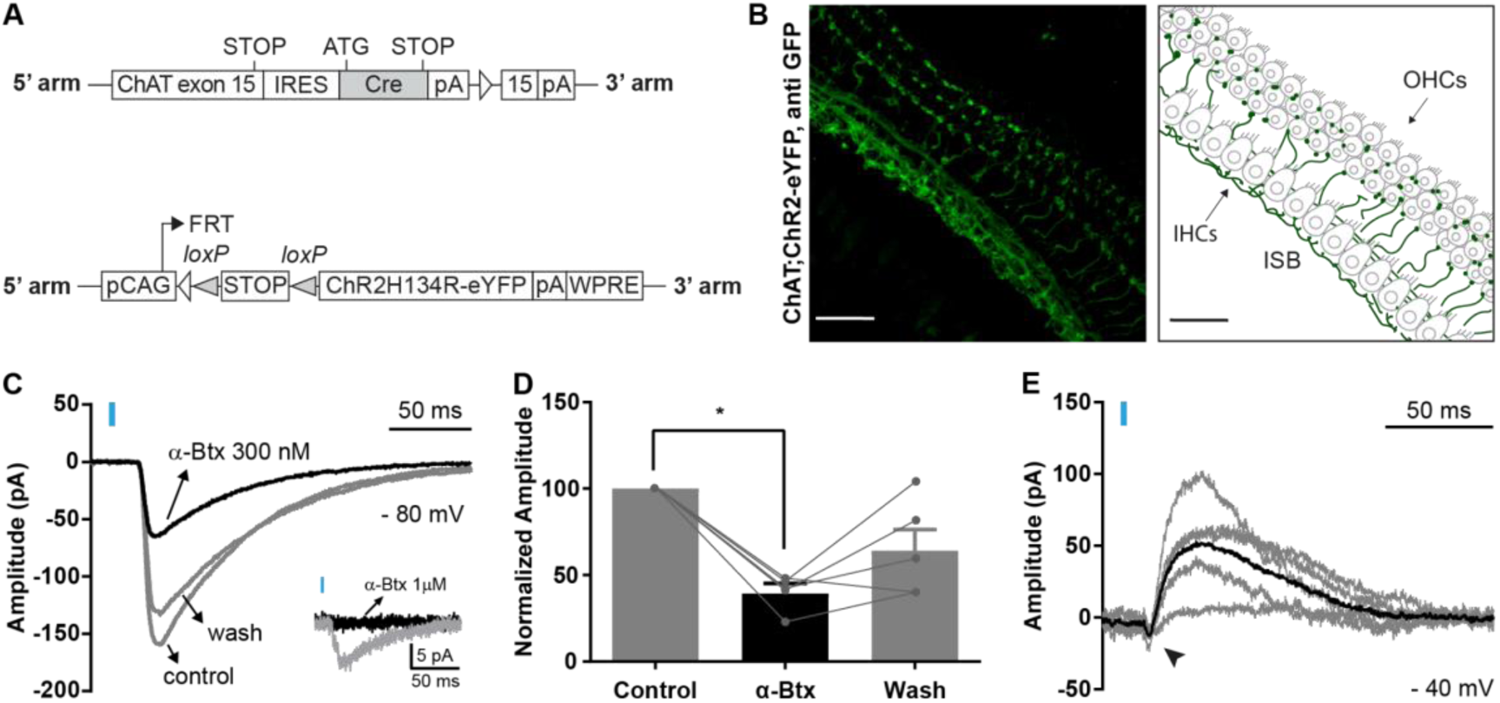
Optogenetic stimulation of cochlear cholinergic fibers. **A.** Schematic representation of the mouse lines transgenes. *Top.* Cre recombinase DNA sequence is inserted within the ChAT gene, restricting its expression to cholinergic fibers. Adapted from Rossi et al. 2011(Rossi et al., 2011). *Bottom.* ChR2-eYFP DNA sequence is inserted within the ROSA locus. Cre excision of a premature STOP codon will only occur in cholinergic neurons, allowing ChR2-eYFP expression. **B.** *Left.* In apical turns of the Organ of Corti of ChAT;ChR2-eYFP mice, the ChR2-eYFP signal was detected with a GFP antibody, showing ChR2 expression mostly restricted to the inner spiral bundle (ISB). Some fibers crossing the tunnel of Corti and reaching the outer spiral bundle were also observed. *Right.* Schematic illustration of the localization of the IHCs and OHCs, which are not visible on the left image. **C.** Representative average traces of one cell showing the effect of 300 nM α-Bungarotoxin (α-Btx), a α9α10 nAChR receptor antagonist, on the amplitude of light-evoked (blue rectangle) *o*IPSCs at *V*h= -80 mV. *Inset.* Representative average traces showing a complete block of *o*IPSCs caused by 1µM α-Btx perfusion (black trace) versus control (grey trace). **D.** Bar graph showing that 300 nM α-Btx caused a significant decrease in *o*IPSC amplitude. Results are expressed as a percentage of control responses (n= 5 cells, 5 mice). * p<0.05, Friedman test. **E.** Representative traces of light-evoked (blue rectangle) *o*IPSCs from the same cell at *V*h= -40 mV. The average trace is shown in black. Arrowhead indicates the small Ca^2+^ inward current mediated by the α9α10 nAChR.

At P9-11, when both the number of functionally innervated IHCs and their sensitivity to ACh reach their maximum (Katz, 2004; Roux et al., 2011), optically-evoked inhibitory postsynaptic currents (*o*IPSCs) were successfully triggered by 2 ms blue light (480 nm) pulses at 0.03 Hz. When voltage-clamped at -80 mV (*V*_h_= -80 mV, E_K_∼ -82 mV), *o*IPSCs of ChAT;ChR2-eYFP mice were inward, had an average amplitude of -123.5 ± 19.29 pA, a decay time constant of 39.57 ± 3.29 ms, and a release probability (Pr) of 0.977 ± 0.02 (13-19 cells, 13 mice, 5-10 single stimulations averaged per recording; **Figure 1C**, control). Furthermore, application of α-bungarotoxin partially (300 nM) and completely (1 µM) blocked light-induced synaptic currents, suggesting that efferent synaptic responses were mainly mediated by ACh activation of α9α10 nAChRs (mean ± SEM: α-Btx (300 nM): -39.29 ± 4.45 pA, wash: -64.01 ± 12.32 pA, Friedman test, *p* = 0.04, 5 cells, 5 mice; **Figure 1C, D *and inset***). At *V*_h_= -40 mV, evoked responses were biphasic with an initial small inward current followed by a longer lasting outward current (69.88 ± 29.26 pA, 4 cells, 3 mice, **Figure 1E**). Except for the release probability, which is higher during optogenetic stimulation (0.98 ± 0.02) compared to electrical stimulation (0.8 ± 0.05; Wilcoxon test, *p*= 0.002; 13-19 cells; 13 mice), these results are consistent with previous experiments detailing MOC synapses onto IHC. Thus, optogenetic stimulation in ChAT;ChR2-eYFP mice induces cholinergic postsynaptic responses mediated by the α9α10 nAChR, similar to the high K and the electrically-evoked synaptic currents previously described (Glowatzki and Fuchs, 2000; Goutman et al., 2005; Zorrilla de San Martin et al., 2010; Kearney et al., 2019).

### ACh is released from GABAergic MOC efferent fibers

Previous works have reported the existence of GABA in the ISB in cells with a morphology consistent with efferent neurons (Fex et al., 1986; Eybalin et al., 1988; Maison et al., 2006), along with an inhibitory effect of GABA on ACh release at the MOC-IHC synapse through presynaptic GABA_B_-R (Wedemeyer et al., 2013). We hypothesized that GABA might be co-released with ACh from MOC terminals. Given that we were able to optogenetically stimulate MOC efferents, an experiment was designed to use this technique in order to stimulate only GABAergic fibers. To this end, we obtained a transgenic mouse line wherein tamoxifen-inducible Cre (CreER^T2^) is under the control of the GAD promoter, and crossed it with a ChR2-eYFP mouse line (GAD;ChR2-eYFP, **Figure 2A**).

**Figure 2.**
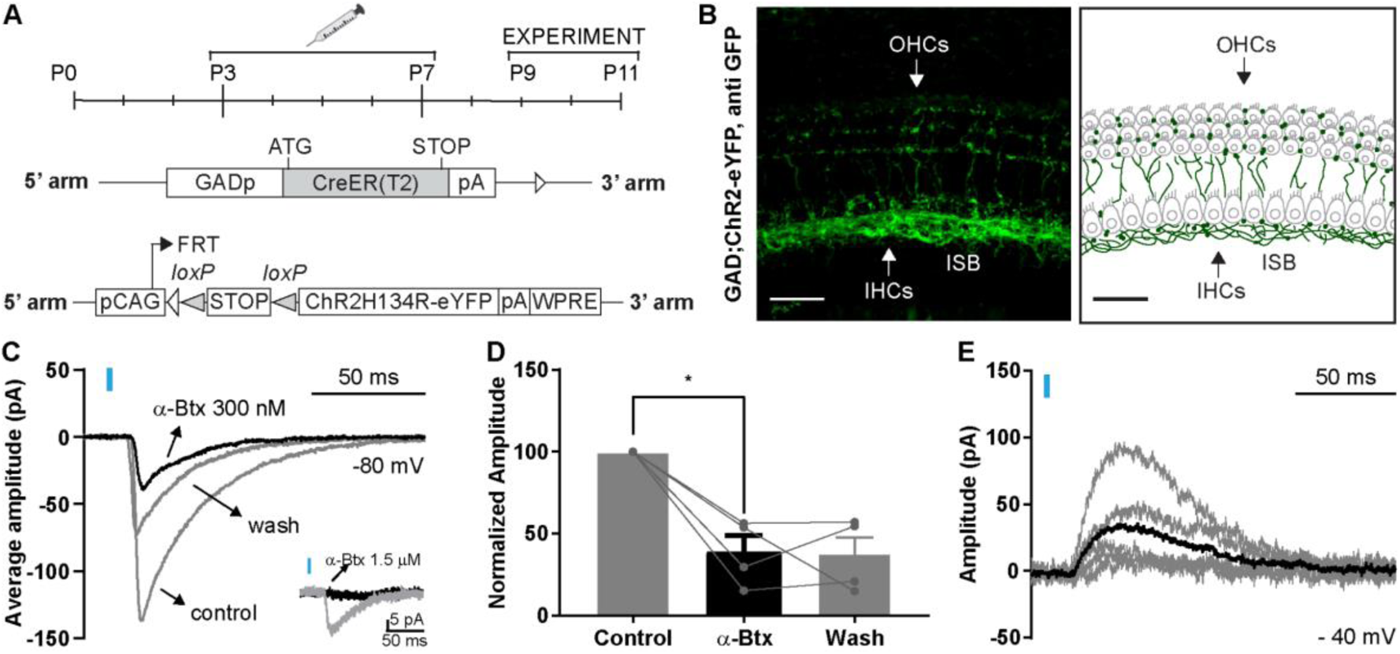
GABAergic fibers projecting to the ISB release ACh. **A.** Schematic representation of the mouse lines transgenes. *Top.* Tamoxifen injection protocol. *Middle.* Cre recombinase is inserted downstream of the GAD promoter, restricting its expression to GABAergic fibers. Adapted from Taniguchi et al. 2011 (Taniguchi et al., 2011). Bottom. ChR2-eYFP is inserted within the ROSA locus. Cre will remove a premature STOP codon allowing ChR2-eYFP expression only in GABAergic neurons. **B.** *Left.* Apical turns of the organ of Corti of GAD;ChR2-eYFP mice labeled for eYFP (with a GFP antibody). ChR2 expression is restricted to the ISB. Some fibers can be seen crossing the tunnel of Corti to reach the outer spiral bundle. *Right.* Schematic illustration of the localization of the neurons relative to IHCs and OHCs, which are not visible on the left image. **C.** Representative average traces of one cell, showing the effect of 300 nM α-Bungarotoxin (α-Btx) on the amplitude of light evoked (blue rectangle) *o*IPSCs at V*h*=-80 mV. *Inset.* Representative average traces show the complete block caused by the perfusion of 1.5 µM α-Btx (black trace) versus control (grey trace). **D.** Bar graph showing that 300 nM α-Btx caused a significant decrease in *o*IPSC mean amplitude ± SEM. Results are expressed as the percentage of the control (n=4 cells, 4 mice).* p<0.05, Friedman test. **E.** Representative traces of *o*IPSC from one cell at V*h*= -40 mV. The average trace is shown in black.

In this mouse model, eYFP fluorescence was found in structures with a morphology consistent with efferent fibers, while the lack of eYFP fluorescence in hair cells suggests that ChR2 expression was limited to efferent terminals (**Figure 2B**). Additionally, using anti-GAD and anti-ChR2 in apical turns of GAD;ChR2-eYFP cochleae (P9-11) we confirmed a significant co-localization of both markers in efferent endings at the base of the IHCs, as well as an expression pattern similar to that observed in the ChAT;ChR2-eYFP mice (Manders’ coefficient = 0.63, **Supplementary Figure 2A**).

Optical stimulation (2-ms light, 480 nm) of GABAergic fibers in GAD;ChR2-eYFP cochlear explants successfully evoked *o*IPSCs that were partially blocked with 300 nM α-Btx (mean ± SEM: α-Btx (300 nM): -38.75 ± 9.89 pA, wash: -36.78 ± 11.03 pA, Friedman test, *p*= 0.04, 4 cells, 4 mice; **Figure 2C, D**) and completely blocked with 1.5 µm α-Btx (**Figure 1C, *inset***). These *o*IPSCs also change polarity at *V*_h_= -40 mV (26.53±7.99 pA, 2 mice, 4 cells; **Figure 2E**) and have amplitude and kinetics similar to those obtained in ChAT;ChR2-eYFP mice and to responses mediated by the α9α10 nAChR (Goutman et al., 2005) (**Supplementary Figure 2B-E**). Importantly, the latency in the onset of the response was not significantly different between GAD;ChR2-eYFP and ChAT;ChR2-eYFP, indicating that the release of ACh is not a result of a disynaptic event (**Supplementary Figure 2E**). Altogether, these results support the notion that the release of ACh occurs during stimulation of GABAergic fibers and strongly suggest that GABA is released from the same MOC terminals.

### GABA is released from synaptic terminals during MOC electrical stimulation

By optogenetically stimulating both cholinergic and GABAergic fibers, we demonstrated that ACh is released from efferent axons and activates nAChRs in the IHCs before the onset of hearing. While presynaptic inhibition of ACh release from MOC terminals by GABA has been reported (Wedemeyer et al., 2013), there is still no evidence confirming that GABA is released from these same cholinergic terminals. Here, we combined the use of the acute organ of Corti preparation and AAV-mediated expression of the GABA sensor iGABASnFR in SGN to gain a better understanding of the source of GABA (**Figure 3**).

**Figure 3.**
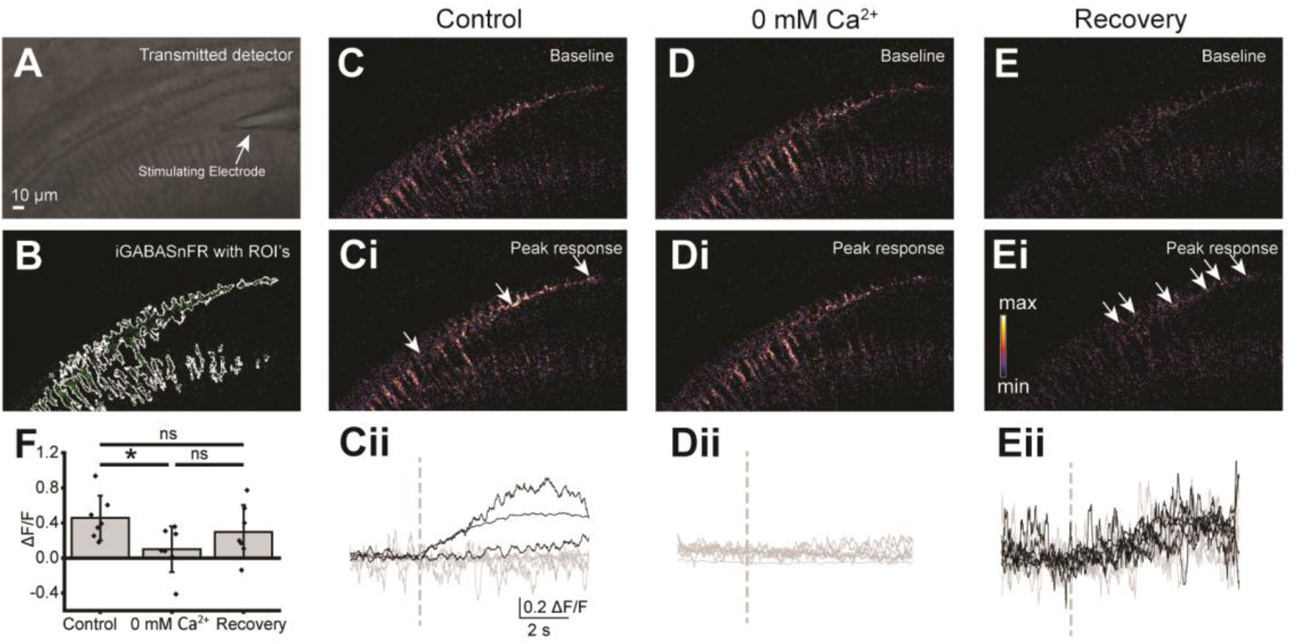
Optical detection of GABA released from efferent neurons onto type I SGN. **A**. Greyscale image of the cochlear region for iGABASnFR imaging in a WT mouse. Stimulating electrode indicated. **B**. Baseline iGABASnFR signal in cochlear neurons. White lines indicate ROIs used for analysis of iGABASnFR fluorescence. Tissue curvature caused by stimulating electrode placement reduces visibility of iGABASnFR fluorescence on the right side of the image. **C**. Baseline iGABASnFR fluorescence in control extracellular solution in an example experiment. **Ci**. iGABASnFR fluorescence in control extracellular solution following electrical stimulation of efferent terminals. Arrows indicate ROIs with a positive iGABASnFR response to electrical stimulation. **Cii**. Rolling window average of timecourse of fluorescence responses of the ROI’s in panels (C-Ci). Black traces indicate ROIs that had a positive response to electrical stimulation, grey traces indicate ROIs that did not respond to electrical stimulation. **D**. Baseline iGABASnFR fluorescence in 0 mM Ca^2+^ extracellular solution. **Di**. iGABASnFR fluorescence in 0 mM Ca^2+^ extracellular solution following electrical stimulation of efferent terminals. **Dii**. Timecourse of fluorescence responses of the ROI’s in panels (D-Di). All ROIs are grey, indicating that no ROIs had a positive response to electrical stimulation in the absence of extracellular calcium **E**. Baseline iGABASnFR fluorescence after recovery to normal 1.3 mM Ca^2+^. **Ei**. iGABASnFR fluorescence in recovery 1.3 mM Ca^2+^ extracellular solution following electrical stimulation of efferent axons. Heatmap scale in inset indicates fluorescence intensity and applies to (C, Ci, D, Di, E, Ei). **Eii**. Rolling window average of timecourse of fluorescence responses of the ROI’s in panels (E-Ei), larger noise due to reduced baseline fluorescence following tissue bleaching. Black traces indicate ROIs that had a positive response to electrical stimulation, grey traces indicate ROIs that did not respond to electrical stimulation. **F.** Quantification of iGABASnFR responses to electrical efferent axon stimulation in type I SGN in control conditions, in 0 mM Ca^2+^ extracellular solution, and after recovery to normal Ca^2+^ extracellular solution. Scale bar in (A) applies to all panels.

The fluorescent GABA indicator iGABASnFR2.0 was transduced in cochlear neurons by injection of AAV into the posterior semi-circular canal of P1-2 mouse pups in WT mice using a serotype and promoter specific for neurons (C57BL/6J: (PHP.eB)-syn.iGABASnFR2-WPRE) or in Bhlhb5-Cre;tdT mice to induce expression specifically in Cre-expressing cochlear neurons ((PHP.eB)-syn.FLEX.iGABASnFR2-WPRE) (Isgrig and Chien, 2018). While expression was lower in Bhlhb5-Cre;tdTomato mice likely because of the required additional step of Cre-mediated recombination, in both mouse lines iGABASnFR expression was limited to cells with the clear morphology of type I SGN and so data was pooled. Cochlear apical turns from injected mice were then acutely dissected for iGABASnFR imaging at P8-11 (6-9 days following AAV injection). Prior to confocal timelapse imaging, a stimulating electrode was placed near the base of IHCs to evoke neurotransmitter release from nearby efferent terminals (**Figure 3A**). iGABASnFR fluorescence (**Figure 3B**) was measured in baseline conditions for ∼3s, then efferent axon stimulation was applied for 1 second (0.26 ms pulse duration, 50 Hz), followed by imaging for ∼6 additional seconds. The fluorescence response (ΔF/F) in each experiment was measured from the peak of the response following axon stimulation divided by the baseline fluorescence prior to axon stimulation (see methods). Electrical stimulation evoked a positive ‘response’ (peak fluorescence greater than the mean of the baseline plus two standard deviations) in a subset of regions of interest (ROIs) that encompassed structures with the clear morphology of type I SGN afferents (mean ΔF/F of positive responses = 0.30 ± 0.25, n = 1-6 ROI’s per cochlea, 14 cochleae from 14 mice). In the subset of experiments in WT mice, iGABASnFR fluorescent responses to efferent stimulation were measured in normal extracellular solution that contains 1.3 mM calcium chloride (control), then again five minutes later in the same region with calcium removed from the extracellular solution (zero calcium) to block calcium influx through VGCC and subsequent neurotransmitter release. Imaging was then repeated in the same location five minutes after returning to control extracellular calcium concentrations (recovery). In these experiments, removal of calcium blocked iGABASnFR responses to electrical stimulation (control ΔF/F = 0.458 ± 0.254; 0 mM Ca^2+^ ΔF/F = 0.101 ± 0.262; recovery to normal Ca^2+^ ΔF/F = 0.297 ± 0.307, One-way repeated measures ANOVA p = 0.020, post-hoc Tukey’s test indicates control significantly different from 0 mM Ca^2+^, n = 7 cochleae from 7 WT mice; **Figure 3C-F**). Together, these results indicate that GABA is released from efferent terminals near the IHCs in an activity-dependent mechanism, suggesting vesicular GABA release from efferent terminals.

### GABA is released from cholinergic MOC efferent fibers

We have shown that ACh can directly be released onto IHCs from GABAergic neurons (**Figure 2**) and that GABA is released at the ISB (**Figure 3**). However, it has not yet been proven that this latter neurotransmitter is liberated from cholinergic fibers, which would cross validate the phenomenon of co-transmission in the cochlea.

According to our previous results (Wedemeyer et al., 2013), the release of GABA at the ISB does not trigger a measurable post-synaptic current in the IHCs but instead acts pre-synaptically on MOC terminals contacting the IHCs. The effect of GABA is through GABA_B_R whose activation reduces the amount of ACh released upon MOC fiber stimulation. Blocking GABA_B_Rs with a specific antagonist, such as CGP 36216, significantly increases ACh release (Wedemeyer et al., 2013). In the following set of experiments, ChAT;ChR2-eYFP mice were used to specifically stimulate cholinergic fibers. Once a basal amplitude of light-evoked responses was determined, the GABA_B_ antagonist CGP 36216 (200 μM) was bath-applied while light stimulation continued (0.03 Hz, pulse duration 2 ms). As shown in **Figure 4**, upon CGP application a potentiated response was observed corresponding to a 36.95% increase in the amplitude of the control response (control: 19.70 ± 4.52 pA, CGP: 26.98 ± 4.71 pA, paired Student’s t test, *p* = 0.007, n = 14 cells, 14 mice, **Figure 4B**). Nevertheless, we observed enormous variability in the effect of CGP on individual cells. Whereas in some cells CGP increased the amplitude of the evoked responses by up to 300%, in others, no changes were recorded (see amplitudes of individual cells before and after CGP 36216 incubation; **Figure 4A**).

**Figure 4.**
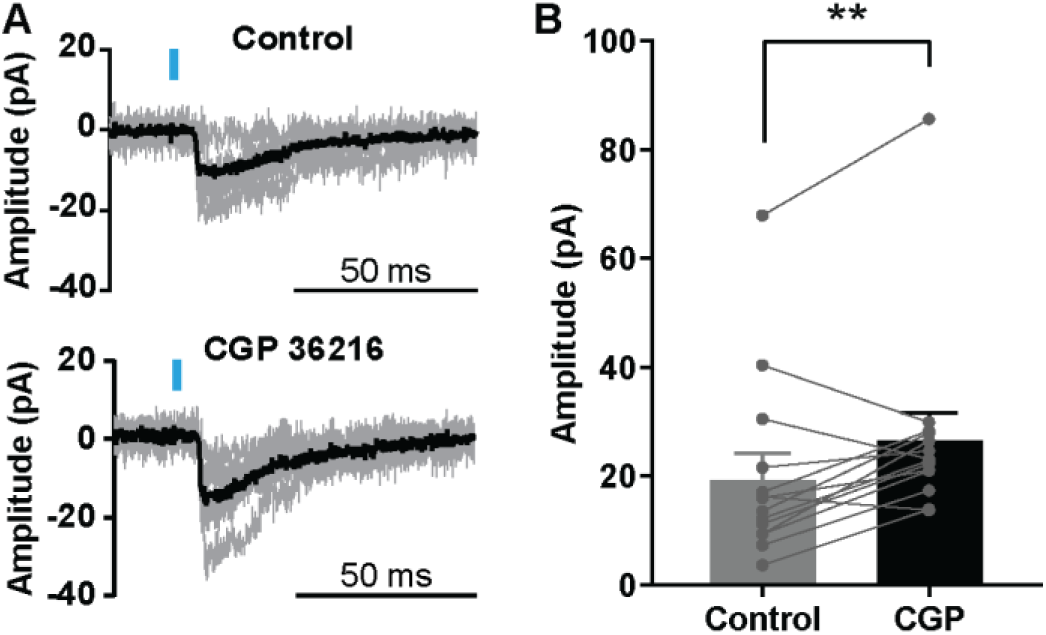
CGP 36216, a specific GABABR antagonist, increases IHC responses to ACh when cholinergic fibers are optogenetically stimulated. **A.** Representative traces of *o*IPSCs from the same cell recorded at a holding potential of Vh=-80 mV before (*top*) and after (*bottom*) 5-7 min incubation with 200 µM CGP 36216. Individual traces are shown in light grey and the average of 5 responses is shown in black. **B.** CGP 36216 (200 µM) caused a significant increase in the amplitude of optogenetically-evoked ACh-mediated currents in ChAT; ChR2-eYFP mice. Individual amplitudes of cells recorded before and after incubation are shown overlaid. Results are expressed as the mean ± SEM (paired Student’s t test, ** p < 0.01, n = 14 cells, 14 mice).

In summary, these experiments indicate that optogenetic stimulation of MOC cholinergic fibers not only caused the release of ACh but also GABA, as the effect of the latter on ACh response amplitude could be prevented by applying the GABA_B_-R blocker (CGP 36216 200 μM).

### Co-localization of cholinergic and GABAergic immunolabeling in efferent neurons

To determine if efferent neurons possess the necessary machinery to synthesize both ACh and GABA, immunostaining experiments were conducted in cochlear whole-mount preparations. The red fluorescence from ChAT;TdTomato mice (P9-P11, apical turn, 2 mice) was used to label cholinergic fibers (amplified with anti-RFP antibody), and an anti-GAD antibody (GAD65) was used to label GABAergic neurons. Co-localization of the two antibodies revealed that there was a subset of efferent terminals that co-expressed cholinergic and GABAergic markers (Manders coefficient = 0.35 ± 0.08) (**Figure 5 Aiii**, **Supplementary Figure 3 B**). However, there were also two populations of terminals that were exclusively either cholinergic or GABAergic.

**Figure 5.**
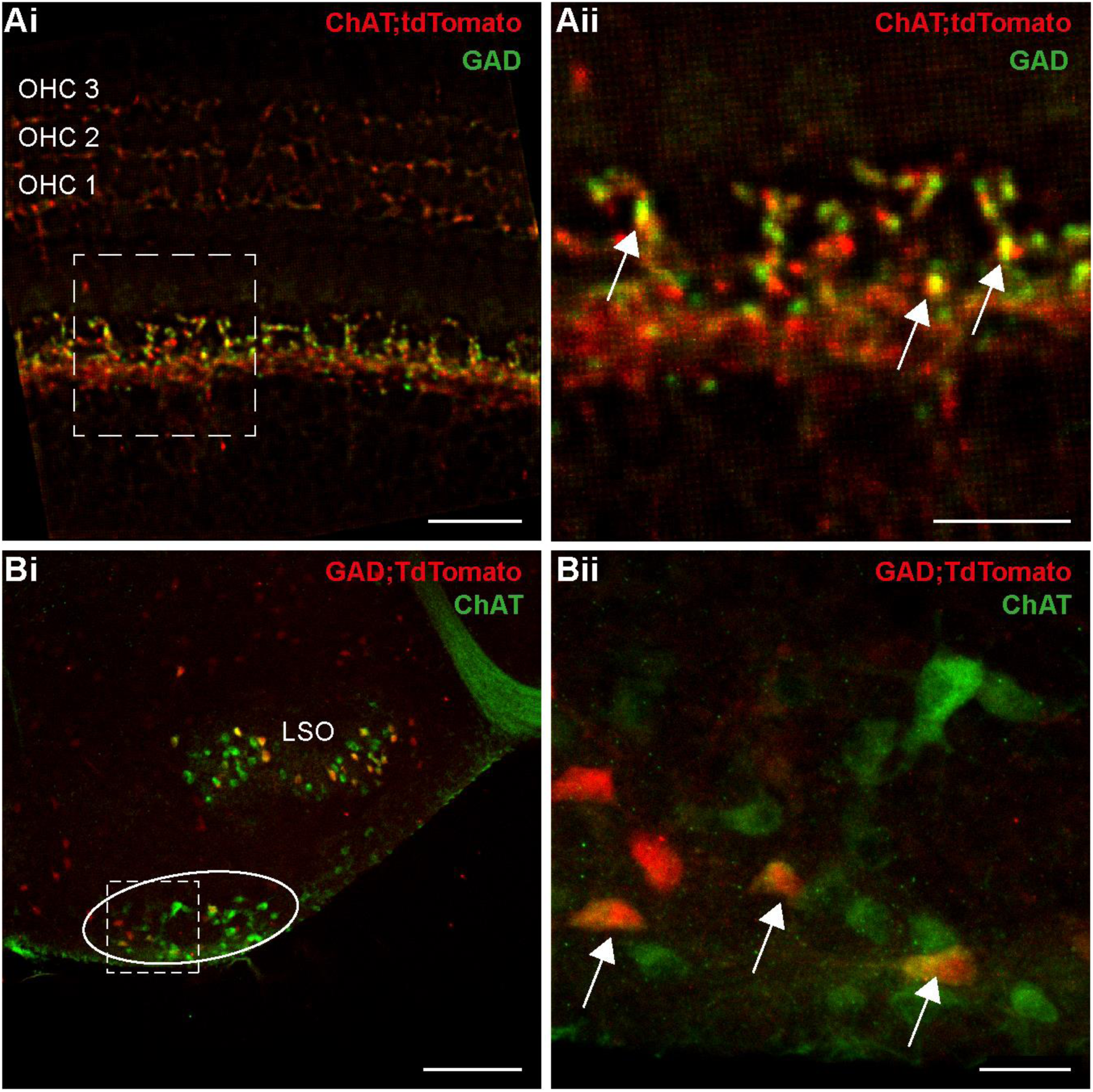
Co-localization of GAD and ChAT antibodies in a subset of efferent terminals and MOC somas. **A.i**. Apical turns of the organ of Corti of ChAT;tdTomato mice (P9-11) were labeled with an antibody against GAD (green). TdTomato fluorescence was enhanced with an anti-RFP antibody (red). Scale bar = 20 µm. **ii.** Higher magnification of the region shown in i (dotted line). White arrows indicate puncta that co-localize. Scale bar = 10 µm. **B.i.** GABAergic labeling in a P9-11 mouse expressing tdTomato under a GAD promoter. TdTomato fluorescence was enhanced with an anti-RFP antibody (red). Anti-ChAT antibody was used to label cholinergic neurons (green). MOC neurons are sparsely localized in the ventral nucleus of the trapezoid body (VNTB, solid line). Scale bar = 200 µm. **ii.** Higher magnification of the region shown in Bi (dotted line). Some neurons show co-localization of both markers (white arrows). Scale bar = 30 µm.

Due to the fact that efferent markers label both LOC and MOC terminals in the developing ISB, immunostaining experiments were also carried out in the brainstem where olivocochlear neurons are spatially segregated (Warr and Guinan, 1979) (**Figure 5 B**). GAD;tdTomato transgenic mice were used to report the presence of GABAergic neurons, which were visualized with an anti-RFP antibody. Cholinergic neurons were labeled with an anti-ChAT antibody. Similarly to the pattern described in the ISB, three distinct neuronal populations were found at the VNTB (18 coronal slices, 6 mice). Out of all the labeled neurons, an average of 54.18% corresponded to cholinergic (11.6 ± 1.5 cells) and 45.81% to GABAergic neurons (13.72 ± 1.5 cells). Additionally, 31.48% of these neurons expressed both markers (7.97 ± 2 cells). These results indicate that a subset of MOC neurons can biosynthesize both ACh and GABA. This supports the evidence obtained from the optogenetics experiments and indicates that both GABA and ACh are co-released from a subset of the same MOC efferent fibers.

### Differential modulation of ACh release sites by GABA

To evaluate the activity of individual synaptic efferent contacts and determine whether they can be modulated by GABA independently from neighboring synapses, we undertook a calcium imaging approach. Given that α9α10 nAChR have a relatively high calcium permeability (Weisstaub et al., 2002; Gomez-Casati et al., 2005), it is possible to measure calcium influx at individual synaptic contacts within an IHC using a fluorescent calcium probe (Moglie et al., 2018). In the experiments in **Figure 6**, the effect of the GABA_B_-R blocker CGP 35348/36216 (100-200 µM) on the MOC-IHCs synaptic activity was evaluated simultaneously both via electrophysiological parameters (average IPSCs amplitude, release probability) and by analyzing calcium indicator responses. As previously shown (Moglie et al., 2018), the advantage of this approach is that multiple synaptic sites onto an individual IHC can be detected and analyzed separately, providing estimates of synaptic parameters for each individual synaptic contact within a given IHC. **Figure 6A** shows representative epifluorescent images taken at the base of an IHC, during the maximum of the postsynaptic current. Representative eIPSCs and fluorescence transients from the same cell are shown in **Figure 6B and C**, both under control conditions and in the presence of CGP. The analysis of 12 cells (12 animals) showed that in the presence of CGP the probability of ACh release increased from 0.53 ± 0.06 to 0.69 ± 0.07 during IHC recordings (*p* < 0.0001, paired t-test, n = 12 cells, 12 mice, **Figure 6E**). **Figure 6D** depicts the amplitude of successful eIPSC events, i.e. excluding synaptic failures, with an average of 99.9 ± 7.8 pA in control conditions and 107.0 ± 11.2 pA in the presence of CGP (*p* = 0.47, paired t-test, ns, n = 12 cells, 12 mice). In the same recordings, a similar analysis was carried out with calcium transients as shown in **Figure 6F and G**, which represent the amplitude of calcium signals and the probability of activation of calcium hotspots per cell, respectively. After the addition of CGP to the bath, a statistically significant increase was observed in the overall probability of activation of calcium hotspots within each IHC (Pr control = 0.11 ± 0.01, Pr CGP = 0.21 ± 0.04, p < 0.014, paired t-Test, n = 12 cells, 12 mice). In addition, no differences were obtained in the amplitude of the calcium signals (ΔF control = 11.8 ± 1.3 A.U., ΔF CGP = 11.6 ± 1.6 A.U., *p* = 0.47, Wilcoxon matched pairs signed rank test, n = 12 cells, 12 mice). Interestingly, the fold change due to CGP effect on the calcium signals was highly proportional to the modulation observed for the synaptic current (**Figure 5H**). These results are compatible with the block of presynaptic GABA_B_-R receptors and therefore of the modulation of ACh release by GABA during MOC neurotransmission (Wedemeyer et al., 2013).

**Figure 6.**
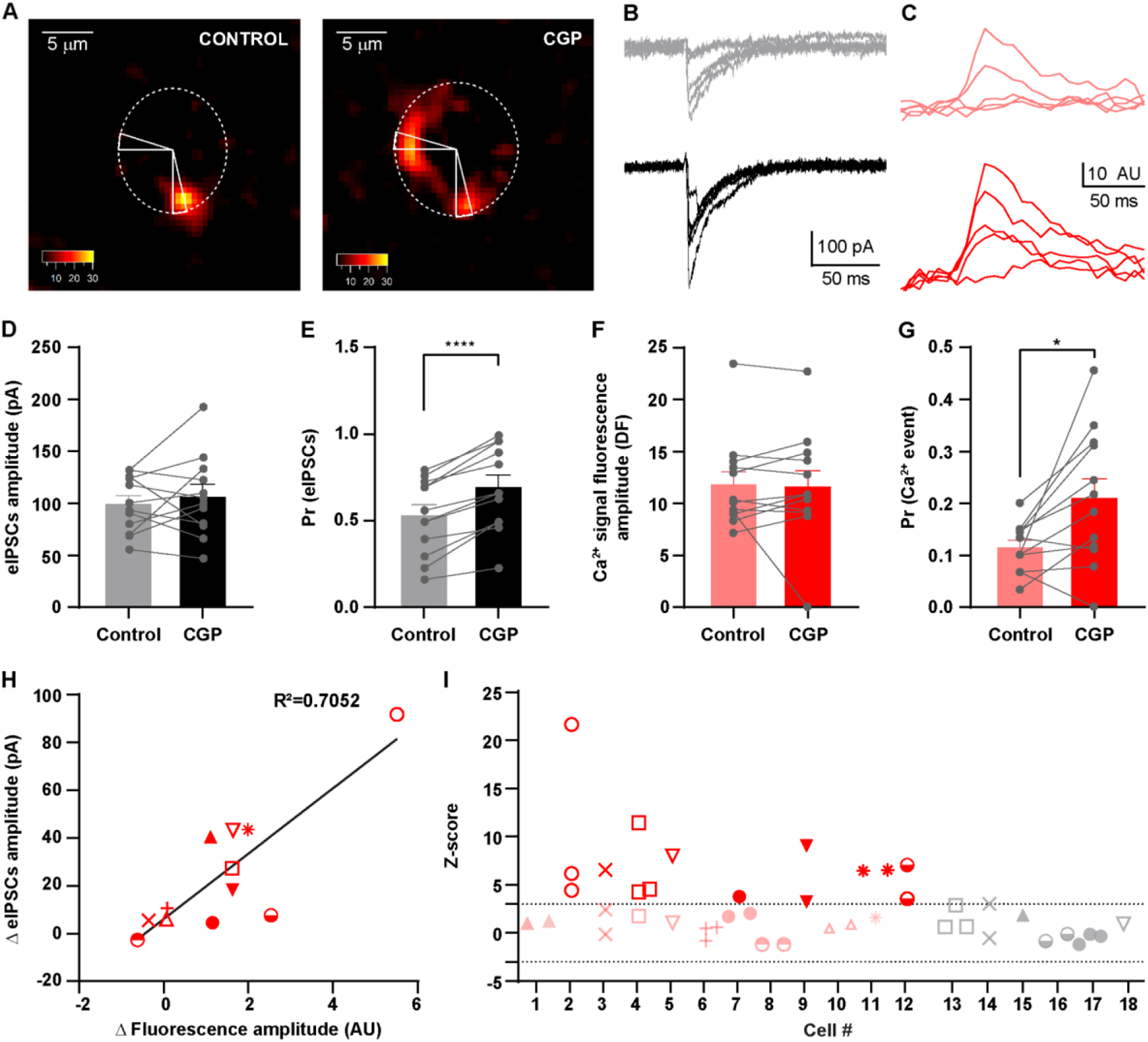
Heterogeneous modulation of MOC efferent terminals by GABA. **A.** Pseudo-colored images of a representative IHC filled with the fluorescent calcium indicator, taken at the peak of the Ca^2+^ transient in response to a MOC stimulation protocol either under control conditions or in the presence of CGP 35348 (100μM). A dotted line indicates the IHC position, and ROIs’ dimensions are delimited by the triangles. **B.** Representative traces of eIPSCs either under control conditions (grey traces, *top*) or with 100 μM CGP (black traces, *bottom*). **C.** Representative traces of calcium signals from one example hotspot in the control experiment (lighter red, *top*) or under the application of 100 μMCGP (darker red, *bottom*). **D.** Average amplitude (without failures) of eIPSCs ± SEM in controls and with 100-200 μM CGP (*p* = 0.47, paired t-test, ns, n = 12 cells, 12 mice). **E.** Probability of release ± SEM in control or with 100-200 μM CGP (**** p < 0.0001, paired t-test, n = 12 cells, 12 mice). **F.** Average amplitude of Ca^2+^ signals (without failures) in control and with 100-200 μM CGP (*p* = 0.47, Wilcoxon test, ns, n=12 cells, 12 mice). **G.** Probability of detecting a calcium event ± SEM in control and with 100-200 μM CGP (*p* = 0.014, paired t-test, * p <0.05, n = 12 cells, 12 mice). **H.** Positive correlation of the effect of CGP in electrophysiological recordings (Δ Amplitude eIPSCs) vs calcium imaging recordings (Δ Amplitude Fluorescence). Symbols represent the same cells used in H, i.e., all the hotspots from the same cell were averaged. **I.** Z-score of hotspot amplitude (with failures) under 100-200 μM CGP (red indicates positively modulated hotspots and pink unaltered hotspots) or control (grey) across different cells. Hotspots from the same cell are indicated with the same symbol. Threshold for a significant effect of CGP = 3.

The average fluorescence intensity increased after MOC stimulation, reflecting a significant increase in the probability of presynaptic ACh release upon CGP application. However, a great heterogeneity in the effect of this drug was found across IHCs and at different hotspots in the same IHC. In order to distinguish between synapses that were modulated by CGP from those that were not (even in the same IHC) in an unbiased manner, we calculated the z-score for each hotspot both in control conditions and in the presence of CGP (see Methods). **Figure 6I** shows the z-score for single calcium hotspots in cells after CGP application (red symbols showing positively modulated hotspots and pink symbols showing unaltered hotspots, 31 hotspots, 12 cells, 12 mice) and in control cells (gray symbols, 12 hotspots, 6 cells, 6 mice). Each symbol type represents individual calcium hotspots from the same cell. The number of hotspots per cell ranged from 1 to 3. Interestingly, a great variation in overall responses were observed across cells: some had hotspots with no statistically significant response to CGP (cells # 1, 6, 8 and 10); in a few, all hotspots were positively modulated by this drug (cells # 2, 9 and 12); and finally other cells presented heterogeneity in the hotspot sensitivity to CGP (some were above and other below threshold, cells # 3, 4, 5, 7 and 11). It is important to note that none of the control hotspots showed a z-score larger than the threshold.

Taken together, these results indicate that there is variability in the effect of CGP on individual synaptic contacts between MOC fibers and IHCs, indicating that ACh release from some terminals is sensitive to GABA modulation, whereas others are not.

## Discussion

In this work, by studying olivocochlear efferent activity onto IHCs before the onset of hearing through the activation of genetically defined neurons, we demonstrate that GABA and ACh are co-released from at least a subset of MOC efferent terminals. Both the pharmacological blockade of cholinergic evoked responses in GAD;ChR2-eYFP mice using specific antagonists of the α9α10 nAChR and the reversal of the current at – 40 mV (**Figure 2**), strongly suggest that optogenetic stimulation of GABAergic MOC fibers in the cochlea leads to the release of ACh. Additionally, optogenetic stimulation of cholinergic neurons resulted not only in the release of ACh but also of GABA, as evidenced by the increase in the amplitude of ACh release after applying a GABA_B_-R antagonist (CGP 36216, **Figure 4**). By using a genetically encoded fluorescent indicator (iGABASnFR) and recording multiple calcium entry sites simultaneously (**Figure 6**), we also demonstrate that GABA’s modulatory effects at the MOC-IHC synapse can vary at different synaptic sites within the same IHC, highlighting the complexity and precise nature of this synaptic regulation.

To enable the co-release of GABA and ACh, the same MOC neurons must express the enzymatic machinery necessary to synthesize both neurotransmitters. This in fact is the case, since immunohistochemical assays performed in this work revealed co-labeling of GAD and ChAT markers in a subset of efferent terminals and MOC somas located in the VNTB (**Figure 5**). These results are in line with those reported in adult guinea pig cochleae where at least half of the efferent fibers visualized at the ISB are immunopositive for GABA, indicating that these GABAergic fibers represent a fraction of the efferent neurons (Altschuler et al., 1984; Fex et al., 1986; Eybalin et al., 1988). In contrast, 100% of cholinergic and GABAergic markers co-localization were reported in MOC-efferent neurons of adult mice (Maison, 2003). This discrepancy can be accounted for by the fact that in our present work we used pre-hearing and not adult mice. NucSeq RNA analysis of MOC neurons in developing mice (P5), have also revealed the presence of transcripts for ChAT, GABA_B_-R 1 and 2 subunits and the GAD2 isoform (Frank et al., 2023). It remains unclear, however, whether these neurotransmitters are co-packaged in the same or in different synaptic vesicles. No evidence to date suggests that GABA is transported by VAChT or that ACh is transported by VGAT, which might indicate that most likely these neurotransmitters are packaged in different vesicular pools, and as a consequence, governed by different release probabilities (Lee et al., 2010; Hnasko and Edwards, 2012; Tritsch et al., 2016).

It is important to note that at P9-11, the developmental stage used in the present study, in addition to MOC fibers that establish axosomatic contacts with the IHCs, LOC fibers also extend to the ISB region (Simmons et al., 1996). Since LOC neurons are both GABAergic and cholinergic (Gulley et al., 1979; Altschuler et al., 1984; Fex and Altschuler, 1986; Eybalin et al., 1988), it is possible that both GABA and ACh are released from these neurons upon optogenetic stimulation of efferent fibers from either ChAT;ChR2-eYFP or GAD;ChR2-eYFP mice. Although unlikely, we cannot rule out the possibility that GABA spillover from LOC neurons ( see Dittman and Regehr, 1997) might contribute to the presynaptic GABA_B_-R-mediated inhibition of ACh release from MOC terminals described in the present work and in Wedemeyer et al. (2013).

Immature IHCs generate spontaneous action potentials (APs) (Kros, 1998; Marcotti et al., 2003; Sendin et al., 2014) during a brief critical period which occurs before the onset of hearing. This spontaneous activity is critical for the correct development of the ascending (afferent) auditory pathway (Kandler et al., 2009; reviewed in Wang and Bergles, 2015). In fact, the absence of a functional efferent system leads to improper maturation of the synapse between the IHCs and the peripheral axons of the spiral ganglion neurons (SGNs) (Johnson et al., 2013). This loss also disrupts the temporal pattern of spontaneous activity in the medial nucleus of the trapezoid body (MNTB), hampering the refinement of its connectivity (Clause et al., 2014; Di Guilmi et al., 2019). Moreover, mice that lack functional postsynaptic α9α10 nAChRs struggle with sound frequency and location processing (Clause et al., 2017). In this scenario, the net inhibition exerted by MOC neurons on the firing pattern of the IHCs becomes highly significant.

At least five mechanisms have been described as potential modulators of the MOC-IHC synapse: 1) a presynaptic negative feedback loop between Ca^2+^ influx through L-type voltage-gated channels and the subsequent activation of BK potassium channels which accelerates repolarization and curtails neurotransmitter release (Zorrilla de San Martin et al., 2010; Kearney et al., 2019), 2) presynaptic GABA_B_-Rs that inhibit P/Q-type voltage-gated channels and reduce the release of ACh from MOC terminals (Wedemeyer et al., 2013), 3) a postsynaptic retrograde messenger, probably nitric oxide, which enhances MOC synaptic transmission (Kong et al., 2013), 4) a presynaptic metabotropic glutamate receptor (mGluR1), likely activated by glutamate spillover from the IHC afferent synapse, that enhances the release of ACh (Ye et al., 2017), and 5) another positive feedback, mediated by released ACh acting through presynaptic nicotinic acetylcholine receptors and causing further release of ACh (Zhang et al., 2020). The interplay between these mechanisms in achieving precise and temporally controlled ACh release remains an open question.

In the case of the presynaptic modulation of ACh release from MOC terminals, a large heterogeneity across different synapses is expected. This is supported by a significant but incomplete co-localization of GABAergic and cholinergic markers in MOC terminals and somata in the VNTB (**Figure 5**). In addition, variability in the net effect of GABA_B_-R antagonists in increasing α9α10 nAChR evoked responses in the IHCs, both with optogenetic (present results) and electrical stimulation (Wedemeyer et al., 2013) is observed. Moreover, calcium imaging experiments (**Figure 6**) indicate that heterogeneity in the effect of the GABA_B_-R antagonist (CGP 36216 or 35348) on the activation of individual calcium hotspots evoked by axon stimulation is due to the differential modulation of ACh release probability at individual MOC terminals.

Heterogeneity in neuromodulatory responses is not an uniqueness of the efferent MOC system. Differential modulation mediated by activity-dependent GABA_B_-R expression has recently been demonstrated in synapses formed by parallel fibers with Purkinje cells in the cerebellum (Orts-Del’Immagine and Pugh, 2018). Additionally, previous research has shown that metabotropic glutamate receptors may be unevenly distributed among boutons of cerebellar parallel fibers, suggesting that variations in spatial distribution can lead to heterogeneous responses (Mateos et al., 1998). Moreover, neuromodulators operate through intracellular second messengers that differentially affect the various types of calcium channels existing in neurons (Brown et al., 2004). At MOC-IHC synapses, both P/Q-and N-type voltage-gated Ca2+ channels (VGCCs) support ACh release (Zorrilla de San Martin et al., 2010; Kearney et al., 2019), however, GABA modulates release by only affecting P/Q-type VGCCs (Wedemeyer et al., 2013). Thus, it can be argued that differences in the sensitivity to GABA might be due to the differential expression of channel subtypes at individual MOC neurons.

The heterogeneity in the expression of GABA markers in MOC neurons could reflect different neuronal subtypes within this system that have been so far neglected. It remains as an intriguing question what role this heterogeneous GABA modulation plays at the MOC-hair cell synapse feedback to prevent excessive ACh-mediated hyperpolarization of the hair cell. Previous studies have indicated that boutons on a given axon can form synapses with different postsynaptic cell types and that much of the heterogeneity in release by presynaptic neurons is due to these different postsynaptic partners (ver intro (Zhang and Linden, 2009). In the case of the cochlea, developing MOC neurons make synaptic contacts with the same cell type (IHCs) and many times presumably with the same individual cell, through local axonal branching (Zachary et al., 2018). Moreover, IHCs are electrotonically compact cells in which an efferent synaptic input at any location produces hyperpolarization throughout (Moglie et al., 2018). Thus, heterogeneity in GABA modulation does not seem to provide any additional integration complexity to the local neuronal circuit. However, one can propose that it is calcium influx through α9α10 nAChR the one that is limited by the GABA negative feedback. This might be important in certain locations of the IHC volume in order to prevent calcium cross-talk with afferent synapses (Moglie et al., 2018). It cannot be precluded that this heterogeneity is a product of a changing synapse at the end of the critical developmental period, right before the onset of hearing in altricial rodents. In other words, a phenomenon that would be more homogeneous at earlier stages of development, becomes more sparse as MOC neurons start to retract from the ISB to extend and reach OHCs.

In summary, our results demonstrate that during the development of the auditory system, GABA is co-released with ACh from a subset of MOC efferent terminals. Through presynaptic GABA_B_-R, GABA exerts heterogeneous modulation at the level of individual synaptic contacts, potentially contributing to the regulation of IHC excitability.

## Author Contributions

Designed research (TC, JDG,MEGC,CJCW,CW), conducted experiments (TC,VCC,SRK,LTC), analyzed data (TC,VCC,SRK,LTC,PIB,VD,JDG, CJCW,CW), wrote first draft of the manuscript (TC, JDG, MEGC,CJCW,CW), wrote manuscript (TC,JDG,CJCW,CW), edited manuscript (TC,JDG,EK, ABE, CJCW,CW), and obtained funding (ABE,EK,CJCW, CW).

## Acknowledgements

This work was supported by Agencia Nacional de Promoción Científica y Tecnológica, Argentina (C.W, A.B.E. and E.K.), and NIH Grant R01 DC001508 (Amanda Lauer, A.B.E.), Intramural Research Program of the NIH, NIDCD, Z01 DC000091 (CJCW)”

The authors declare no competing financial interests.

## Notes

### Competing Interest Statement

The authors have declared no competing interest.

