## Supplementary Figures for "Co-release of GABA and ACh from medial olivocochlear neurons fine tunes cochlear efferent inhibition"

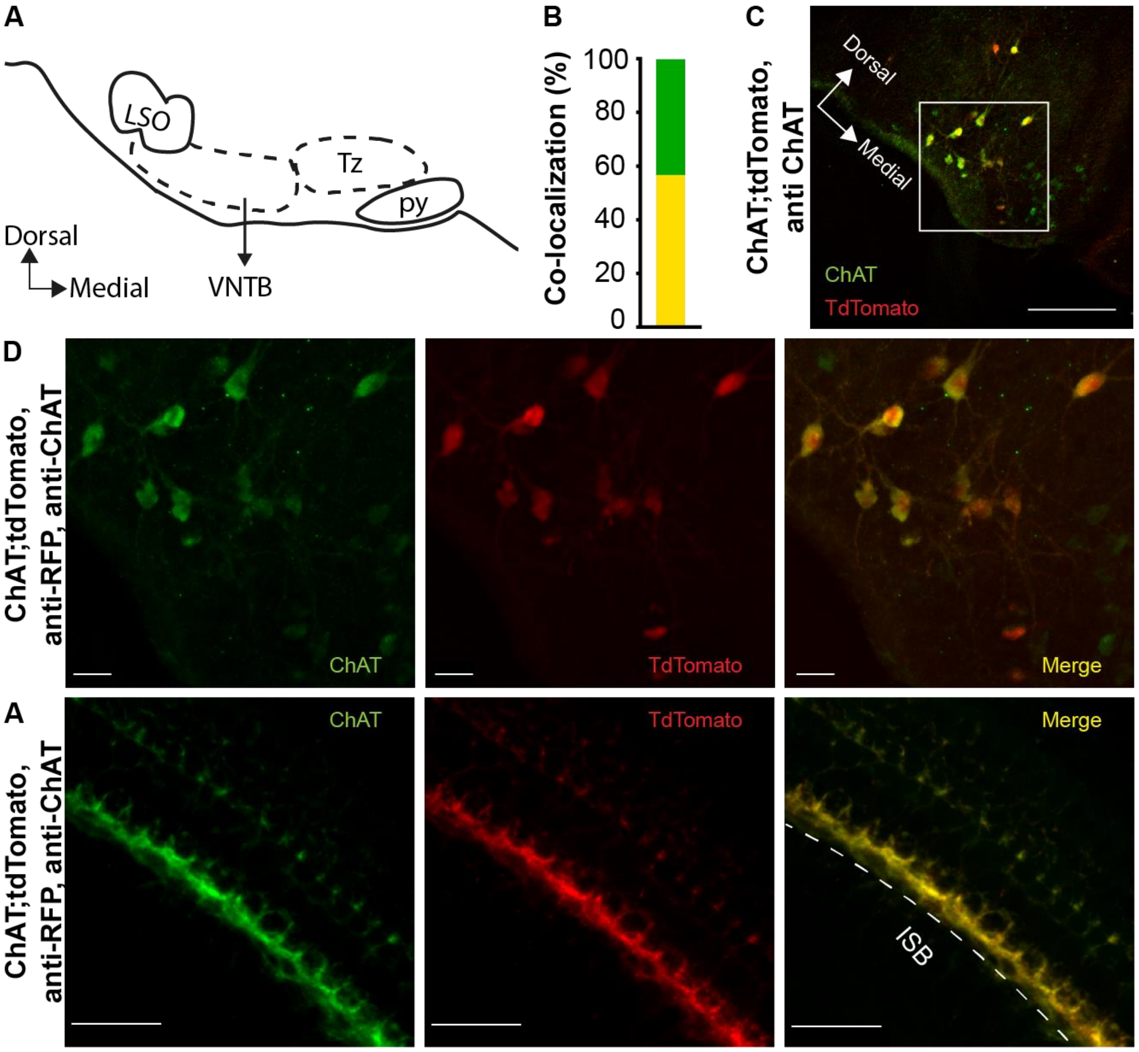


**Supplementary Figure 1. Co-localization of anti-ChAT and tdTomato (anti-RFP) expression in ChAT;tdTomato mice. A.** Schematic diagram of auditory nuclei in the mouse ventral brainstem. LSO: Lateral Superior Olive; VNTB: Ventral Nucleus of the Trapezoid Body; Tz: Trapezoid Body; py: Pyramidal tract. Adapted from Paxinos, George, and Keith B.J. Franklin . The mouse brain in stereotaxic coordinates: hardcover edition. Access Online via Elsevier, 2001. **B.** Colocalization percentage of ChAT and tdTomato in MOC somas located in the VNTB. An anti-RFP antibody was used to enhance tdTomato fluorescence. Yellow: anti-ChAT+;anti-RFP+ cell count (56.6%), Green: anti-ChAT+;anti-RFP- cell count (43.4%). **C.** Immunolabeling against ChAT (green) and RFP (red) in the brainstem of ChAT;tdTomato mice. Scale bar = 200 mm. **D.** Higher magnification of the area shown in C (white square). Scale bar = 30 mm. **E.** Apical turns of the organ of Corti of ChAT;tdTomato mice with labeled ChAT (left) and RFP (middle) antibodies. The co-localization of both markers can be seen in the ISB and near OHCs (merge, right panel). Scale bar = 30 mm.

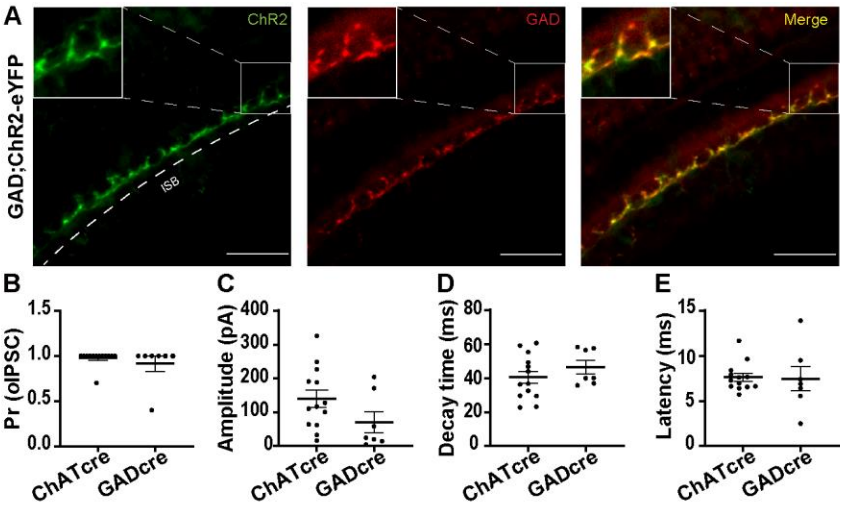


**Supplementary Figure 2**. **Synaptic properties of light-evoked IPSCs in GAD-Cre vs ChAT-Cre mice. A.** Apical turns of the organ of Corti of P9-11 GAD;ChR2-eYFP mice were labeled for eYFP (anti-GFP antibody, green, left panel) and GAD (anti-GAD65 antibody, red, middle panel). Expression is restricted to the inner spiral bundle (ISB, dashed line) and both markers partially co-localize (Merge, yellow, right panel). Scale bar = 30 µm. **B, C, D and E.** Plots of release probability (B, ChAT-Cre: 0.98 ± 0.02, GAD-Cre: 0.91 ± 0.08), amplitude (C, ChAT-Cre: 139.8 ± 25.41 pA, GAD-Cre: 71.37 ± 31.05 pA), decay time (D, ChAT-Cre: 40.54 ± 3.63 ms, GAD-Cre: 46.55 ± 3.95 ms), latency (E, ChAT-Cre: 7.62 ± 0.44 ms, GAD-Cre: 7.45 ± 1.34 ms) in GAD;ChR2-eYFP and ChAT;ChR2-eYFP *o*IPSCs. The mean and SEM are shown. Dots correspond to individual values from one cell. No significant differences were found in all cases (Mann Whitney test, ChAT N=13, GAD N=7).

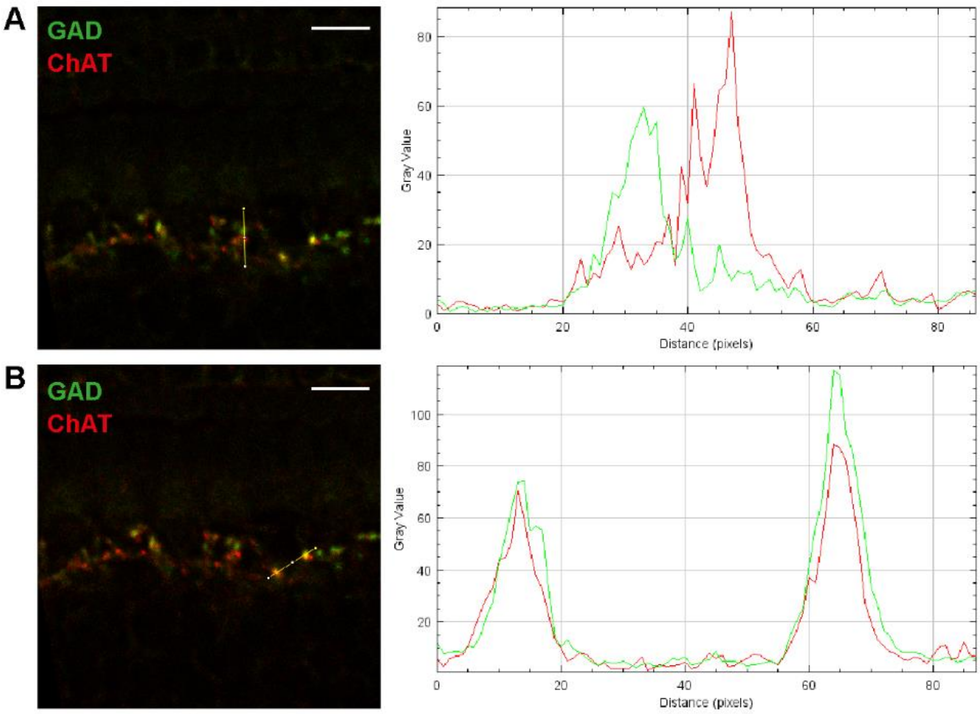


**Supplementary Figure 3. Cholinergic, GABAergic and Cholinergic/GABAergic terminals within the ISB. A.** *Left.* Immunolabeling against GAD and tdTomato in the apical portion of the cochlea of ChAT;tdTomato mice. Different lines (yellow) were drawn in just one confocal image to plot the pixel intensities of both channels shown on the right. *Right.* Plot profile of the pixel intensities in both channels, showing solely GABAergic (green) and cholinergic (red) puncta. Scale bar = 10 µm. **B.** Same as in A, but the line profile shows two puncta that colocalize.
